# Scalable Genotyping of Microbial Colonies

**DOI:** 10.1101/2023.12.06.570470

**Authors:** Arnold Chen, Nkazi Nchinda, Nate Cira

## Abstract

The sequence of the 16S region is taxonomically informative and widely used for genotyping microbes. While it is easy and inexpensive to genotype several isolates by Sanger sequencing the 16S region, this method becomes quite costly if scaled to many isolates. High throughput sequencing provides one potential avenue for obtaining 16S sequences at scale, but presents additional challenges. First, DNA purification workflows for high-throughput sample preparation are labor intensive and expensive. Second, cost-effective multiplexing and library preparation schemes are difficult to implement for many libraries on a single sequencing run. Therefore, we implemented a scalable protocol for isolate genotyping involving colony polymerase chain reaction (PCR) with simple cell lysis as well as a four barcode indexing scheme that enables scalable multiplexing and streamlined library preparation by amplifying with four primers simultaneously in a single reaction. We tested this protocol on 93 colonies cultured from environmental samples, and we were able to ascertain the identity of ∼90% of microbial isolates.

## INTRODUCTION

Growing microbes in isolation allows them to be characterized and utilized. Libraries of microbial isolates are collected for many purposes including: production of natural product metabolites for drug discovery (1), recapitulating representative microbial communities such as gut microbiomes (2), investigating organism ecology and evolution (3), cataloging clinical isolates (4), and the growth and discovery of new microbes (5). Frequently isolates from such libraries need to be genotyped to, for example, identify organisms of interest (6), dereplicate clones (1), identify which clones have a gene of interest (7), and determine organism relatedness (3).

Genotyping a single gene for a single isolate is straightforward, commonly accomplished with Sanger sequencing. Starting with a grown isolate, this process often involves microbial lysis, DNA purification, DNA amplification, and product cleanup before Sanger sequencing and analysis. This can be accomplished at reasonable costs for a few isolates, but quickly becomes time-consuming and costly when scaling to large collections of isolates.

High-throughput sequencing generates far more sequenced bases per sequencing run than Sanger sequencing, and is widely used to analyze many organisms at once in the study of microbial communities (8-10). However, each sample must be prepared and made into a library independently, which is typically even more burdensome than preparing samples for Sanger sequencing. For a large collection of isolates, library production becomes a significant barrier.

Therefore, we developed a workflow for streamlined production of high-throughput amplicon sequencing libraries. We show how this workflow enables scalable genotyping by sequencing the 16S V4 region of a library of microbial isolates.

## MATERIALS AND METHODS

### Overview

Here we describe the scalable genotyping protocol (Fig. 1A) we devised which features 1) colony polymerase chain reaction (PCR) with simple lysis, 2) indexing conducive to scalable multiplexing across multiple well plates, and 3) four-barcode indexing performed in one reaction, excluding the need for intermediate cleanups and additional reactions. In this section, we describe our protocol in detail, referencing past work that inspired each step. In addition, we discuss the rationale behind our specific approach and the tradeoffs compared to alternative methods.

**Figure 1.**
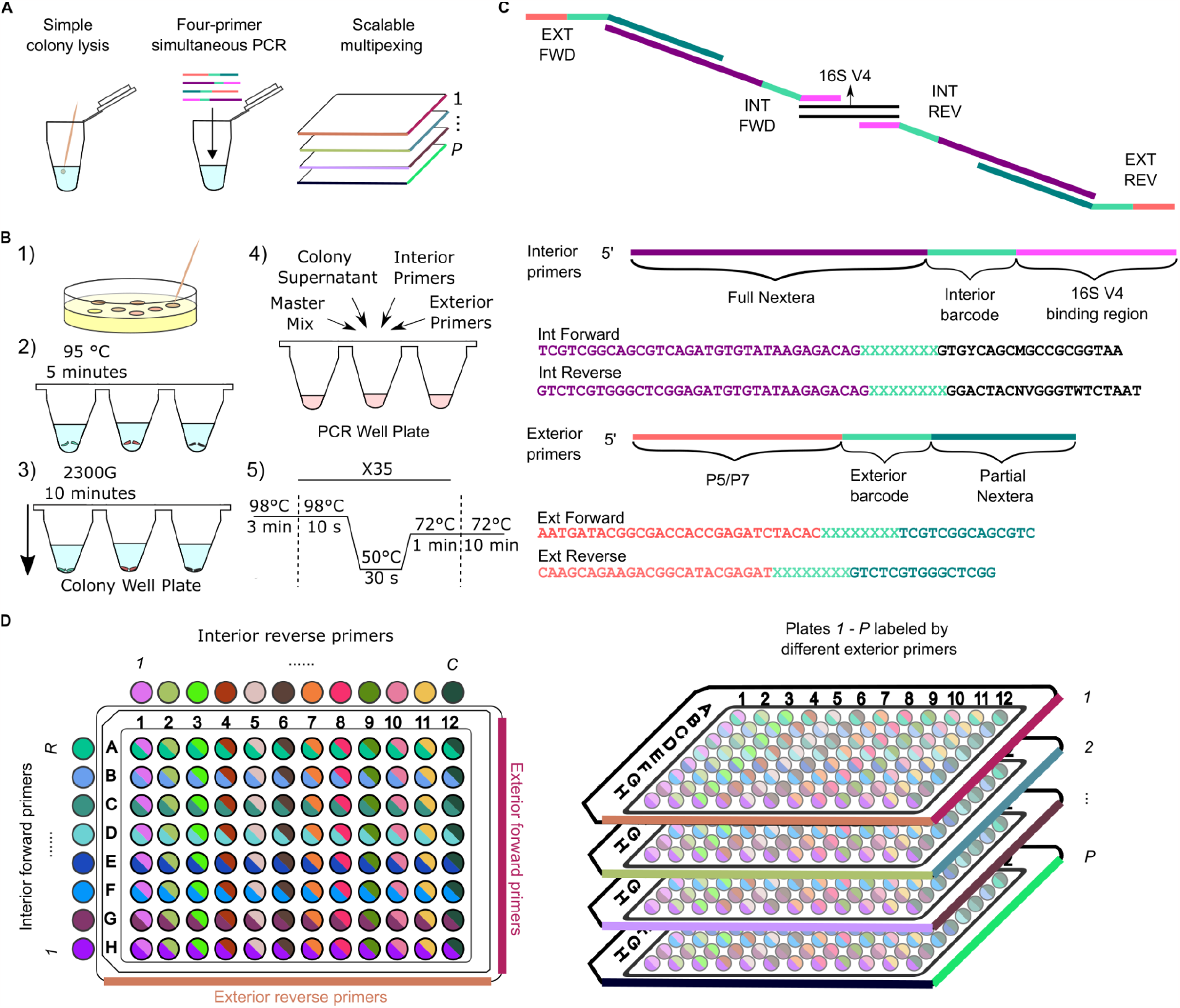
a) An overview of the features of our protocol b) Each colony was picked from agar plates, placed in water, heated, then centrifuged. The supernatant was extracted and mixed into PCR reaction mix containing Master Mix and both interior and exterior primers. The reaction was thermocycled to arrive at the final amplicon. c) Interior primers anneal to the 16S rRNA V4 region, and exterior primers bind to the Illumina flow cell. The final amplicon contains both interior and exterior sequences due to a shared Nextera region between the interior and exterior primers. Exterior and interior primers have both forward and reverse sequences, and each sequence encodes a unique index. The final amplicon contains four different indices. d) The library indexing scheme encodes rows with interior forward primers and columns with interior reverse primers. One set of exterior primers was used throughout the whole well plate (left). The scheme is easily scalable to multiple well plates (right) by using exterior indices to label each plate.

### Sample collection, colony picking, and colony freezer stocks

10 environmental samples were collected from various locations near Cornell University, Ithaca, NY. Each sample was prepared with different dilution treatments including undiluted, 10^−1^, 10^−2^, and 10^−3^ dilutions then plated on tryptic soy agar (TSA), 5% sheep blood in tryptic soy agar, and Luria agar (LA) plates. Plates were incubated at room temperature for three days.

93 colonies were selected, with 31 colonies from each type of agar plate, being careful to exclude colonies that were in contact with another colony. In addition, freezer stocks of the selected colonies were prepared for future cataloging.

### Colony lysis

The first step of most genotyping workflows is to prepare purified DNA from the sample, often using commercial kits or protocols that become costly and onerous to perform at a large scale. A more scalable approach is colony PCR (11), which involves processing microbial colonies without rigorous purification, instead incorporating only simple procedures such as heating and centrifuging before downstream reactions.

Here we adapted a colony PCR protocol (12). Colony material was placed into wells of a 96 microwell plate filled with 80 μl of PCR-grade water (Invitrogen) (Fig. 1B). The microwell plate was sealed with PCR plate foil, heated at 95 °C for five minutes to crudely lyse the colonies, and centrifuged at 2300g for 10 minutes to pellet the cellular debris. The supernatant of each colony was used as DNA template for the PCR. This approach, while unlikely to provide a high concentration of purified template DNA, avoids the costs and time associated with more involved sample preparation methods.

### Primer design

A single high-throughput sequencing run generates ample sequencing reads, allowing cost-effective pooling and sequencing of numerous samples simultaneously. When pooling samples, a multiplexing scheme must be used to determine which sample each read originates from. Multiplexing is accomplished by introducing different indices into each sequencing library to encode the sample of origin. A simple scheme is to incorporate a different index for each sample. This requires ordering *N* primers with unique indices for *N* samples.

Alternatively, combinatorial dual indexing can be used, where indices are repeated within the same row and column in a well plate format, and each sample is encoded by a unique row/column combination. This reduces the required number of primers to 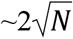 However, this indexing scheme is susceptible to an undesirable phenomenon called index hopping (13), where indices are misassigned on Illumina sequencers. Since libraries in the same row or column differ by only the forward or reverse index, misassigning either index will lead to erroneous identification of the sample.

To address this, a unique dual indexing scheme can be used, where each library uses two unique indices, requiring 2*N* total primers. If either the forward or reverse index is misassigned, this error can be inferred from the other index. However, the large number of unique primers in this scheme becomes prohibitively expensive (Fig. 2A).

**Figure 2.**
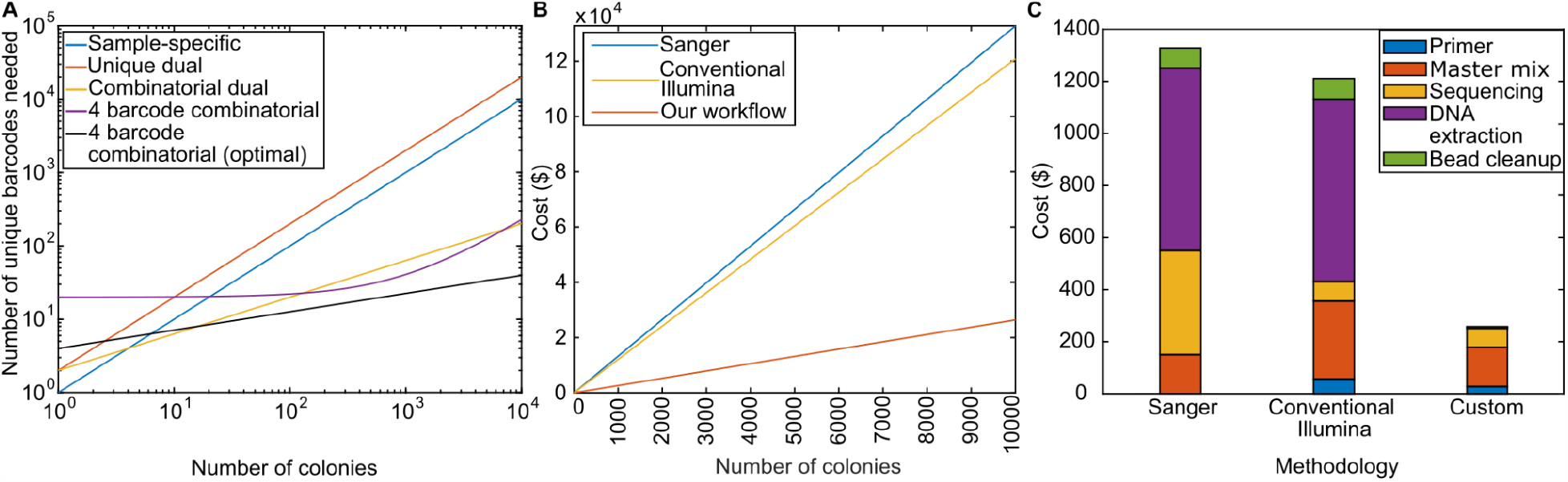
a) Number of unique primer barcodes needed for a given number of colonies using 96 well plate formats. The sample-specific and unique dual indexing schemes scale linearly with colonies. The four barcode combinatorial indexing scheme using the exterior barcodes as dual unique plate labels scales linearly with colonies for larger numbers of colonies but requires fewer barcodes than the sample-specific and unique dual index schemes. It is comparable to the combinatorial dual scheme in efficiency but does not suffer from index hopping. The optimal four barcode combinatorial indexing scheme scales very efficiently, with the fourth root of the number of colonies. b) The cost of different microbial genotyping workflows comparing Sanger, Illumina, and our custom workflow. c) The estimated cost breakdown of different genotyping workflows per 100 isolates. See SI for a detailed table of costs and explanations.

To enable scalable multiplexing, we opted for a four barcode multiplexing scheme using four primers. The scheme includes exterior forward, exterior reverse, interior forward, and interior reverse primers (Fig. 1C). The primers are referred to as “exterior” and “interior” because in the final amplicon, the interior primer sequence is situated in between the target DNA of interest and exterior primer sequence. Interior primers anneal to the target DNA of interest while exterior primers contain the Illumina P5/P7 regions that allow binding to the flow cell. The exterior primers overlap with the outermost region of the product created by the interior primers, resulting in a longer amplicon with all the necessary sequence regions for indexing and sequencing. To allow for multiplexing, we incorporated 8 base pair (bp) barcode sequences to all four primers.

To target the ∼254 bp 16S V4 region, we incorporated primer sequences 505F (14) and 806R (15) for the forward and reverse interior primers, respectively. The target overlapping regions on these primers are easily modifiable to target other regions while still maintaining the four barcode multiplexing capabilities.

We used four indices for each amplicon, using interior indices to label the well location and exterior indices to label the plate. We barcoded a 96 microwell plate with unique pairwise combinations using eight different interior forward indices along the rows and twelve different interior reverse indices along the columns (Fig. 1D). One exterior forward and one exterior reverse index was used to label the plate. This version of a four barcode scheme requires ∼(2*P* + *R* + *C*) unique primers, where *P* is the number of plates and *R* and *C* are the number of rows and columns in each plate. Across multiple 96-well plates, this requires 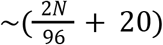 total primers, which scales better than the sample-specific and unique dual indexing schemes.

Theoretically, this scheme can be very primer-efficient, allowing up to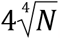, though this would often require inconvenient index layouts with respect to processing in well plates.

### Library preparation and amplicon sequencing

Prior work has demonstrated a sample-specific one barcode scheme involving four primers through sequential amplification (16). A four barcode scheme involving four primers typically requires sequential addition of reagents and multiple amplification steps (17-19). Other work has shown simultaneous addition of reagents and amplification of four primers through indexing of dual unique barcodes (20) or dual identical barcodes (21). In our approach, we carried out a four barcode scheme where we simultaneously added all four primers which contained all the necessary indices and adaptors into the reaction mix under one thermal cycled reaction. Our approach omits the need for additional reactions, thus reducing costs and avoiding the parallel cleanup between different rounds of amplification. This allows us to reap the benefits of scaled multiplexing without the effort of additional reactions.

Each 25 μL reaction volume contains 2 μL of colony supernatant, 0.05 μM of each interior primer, 0.45 μM of each exterior primer, and 12.5 μL of master mix (2X Phusion High-Fidelity PCR Master Mix, New England Biolabs). The high ratio of exterior to interior primers helps ensure the dominant product is the desired full-length amplicon. The presence of some interior amplicon is not problematic, as it will not sequence without illumina adapters. The reaction was thermal cycled with the following conditions: initial denaturation at 98 °C for three minutes, followed by 35 cycles of denaturation (98 °C for 10 s), annealing (50 °C for 30 s), and extension (72 °C for 60 s) and a final extension step at 72 °C for 10 min.

PCR products were pooled then purified with AMPure XP beads (Beckman Coulter) at 0.7x bead/sample ratio. Sequencing was performed with the Illumina MiSeq v3 kit to obtain approximately 800,000 300 bp paired-end reads at Cornell Institute of Biotechnology (Ithaca, NY).

### Bioinformatic analysis

Raw reads were demultiplexed by the interior indices to assign well location. DADA2 was used to filter the reads using default parameters. Excluding the barcode and binding regions, forward reads were further truncated to 180 bp, while reverse reads were truncated to 120 bp, based on an initial assessment of per base sequence quality. Reads were merged to create amplicon sequencing variants (ASVs). ASVs were deemed representative of a well if the ASV read abundance in the well passed a minimum read threshold of 15 reads and a minimum proportion of 0.1 of the total read abundance of that well. We determined the taxonomy associated with each ASV by aligning reads with the SILVA database using VSEARCH. A phylogenetic tree was created by first performing multiple sequence alignment through MUSCLE then using MEGA11 to construct maximum likelihood trees with 100 bootstrap replicates.

### Sanger sequencing and analysis

Freezer stocks corresponding to wells with two or more representative ASVs were plated and grown on agar plates of the corresponding media type. The colonies were purified (Qiagen DNeasy Powersoil Pro kit). Amplification and product cleanup were performed. The products were submitted for Sanger sequencing at the Cornell Institute of Biotechnology. The Sanger chromatograms were examined and sequences were aligned with ASVs to identify matches.

## RESULTS AND DISCUSSION

To assess the performance of our workflow, we mapped the representative ASVs to well positions (Fig. 3A). Overall, 96% of the wells amplified successfully and had at least one representative ASV (Fig. 3B). We taxonomically identified these organisms from environmental samples, finding organisms from two phyla: Firmicutes and Proteobacteria. There were eight unique genera, of which three were gram-positive: *Exiguobacterium, Bacillus, Lysinibacillus*, and five were gram-negative: *Pseudomonas, Acinetobacter, Shewanella, Serratia*, and *Aeromonas* (Fig. 3C).

**Figure 3.**
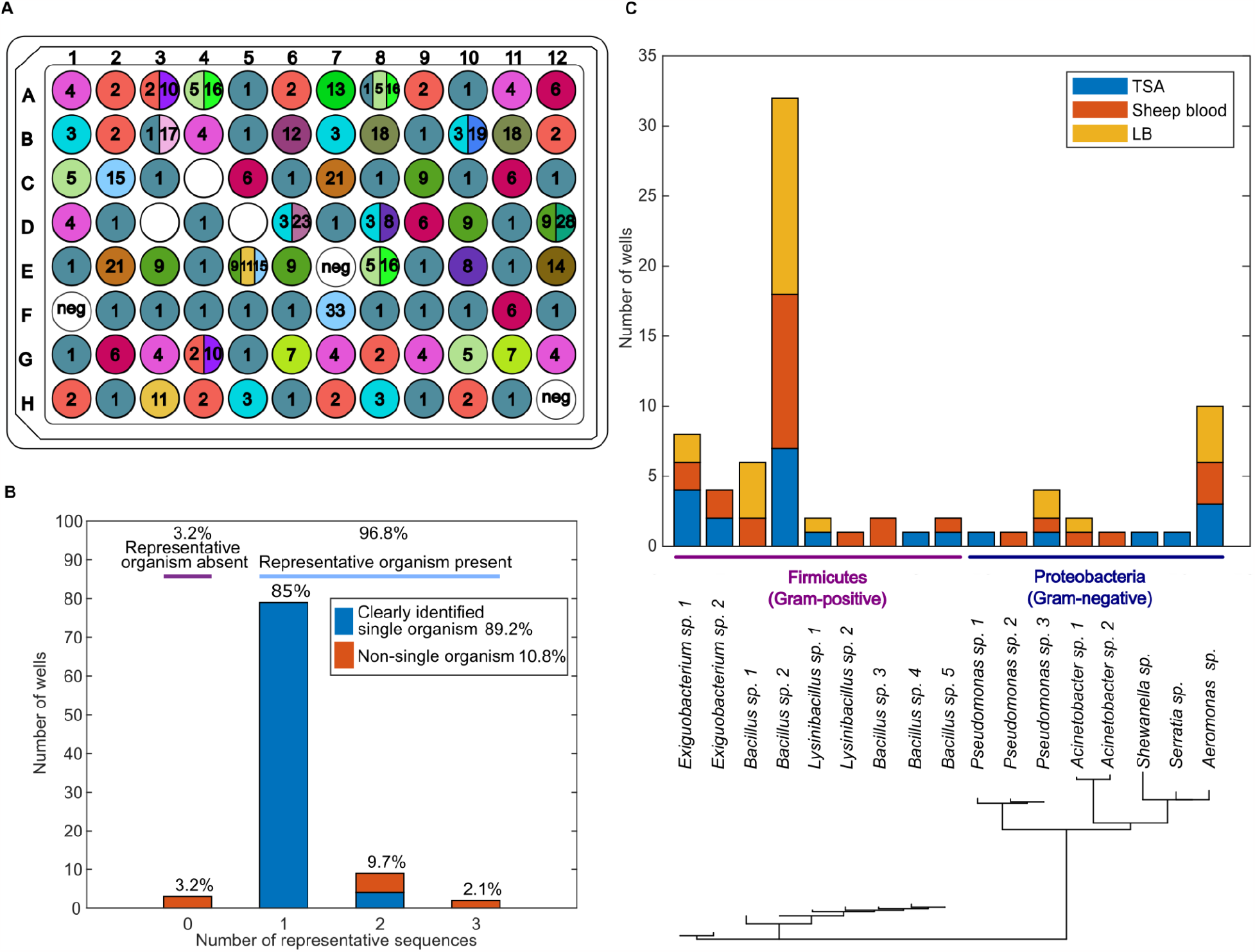
a) Representative ASVs in each well. Each number and color indicate a unique ASV. White wells indicate no representative ASV was detected. b) The proportion of wells that matched to between 0-3 representative ASVs, colored and labeled with the proportion of wells that contained representative organisms and the proportion of wells where we were able to clearly identify the organism’s taxonomy. c) The agar plate media that each organism was represented in and the associated phylogenetic tree.

Next, we categorized the wells based on the number of ASVs they mapped to and further investigated select wells through Sanger sequencing. Out of 96 wells, the three negative controls did not yield any representative ASVs. Of the 93 wells with isolates, 79 wells matched to only one representative ASV, providing clear taxonomic identification of those organisms. For confirmation, we replated and Sanger sequenced freezer stocks from four of these wells which indeed matched with the original ASV sequences. In the remaining 17 wells, 11 mapped to more than one representative ASV (Fig. 2B). We investigated the identities of organisms in these 11 wells through regrowth of the freezer stock and Sanger sequencing. We obtained sequences from eight out of the 11 wells, but poor amplification and failed Sanger sequencing prevented obtaining sequences from the remaining three wells. Out of the eight sequences, all sequences matched to one of the ASVs in the corresponding wells (Supplementary Table S1), supporting the validity of the ASVs and hinting that the presence of multiple ASVs may be attributed to mixed colonies. In four out of these eight wells, the multiple ASVs within a well only differed by one base pair, which was also observed in Sanger sequencing as a mixed signal at the same base pair discrepancy (wells A3, A4, B10, E8) (Supplementary Figure S1). This may be attributed to organisms possessing intragenomic variation which is well documented for the 16S rRNA gene (22, 23). Three wells contained non-amplified organisms, potentially due to fungal colonies without a 16S region, PCR inhibitors, or the colony’s susceptibility to lysis (24, 25). These results demonstrate the utility of this workflow to properly identify the taxonomy of a collection of microbial isolates. Overall, 90 out of 93 wells (96.8%) yielded representative sequences, and 83 out of 93 wells (89.2%) indicated a single clear organism.

In addition to successful identification of isolates, cost is a key aspect to consider for a scalable workflow. We compared the cost of three workflows: Sanger sequencing, conventional Illumina sequencing with two-step PCR, and our custom approach (Fig. 2B). Note that we consider the costs on a per-colony basis. In certain cases, there may be an initial upfront cost involved with obtaining the required reagent stock at a volume larger than required for the experiment. However, the excess reagent stock can often be used in future experiments, and average cost calculation becomes especially applicable in scaled-up experiments.

In all three workflows, the cost scales linearly with the number of colonies under the above assumptions. Sanger sequencing has the highest per-colony cost, followed closely by conventional Illumina sequencing. Our custom workflow has less than half the per-isolate cost compared to the other methods.

The contribution to the total cost comes from various factors including DNA preparation, reagents (primers and master mix), sequencing, and purification of PCR products (Fig. 2C) (Supplementary Table S2). One key factor that led to the reduced cost of our scheme is the omission of DNA prep, which accounts for a major fraction of the cost in the other two schemes. Another benefit is that only one bead purification is required for our scheme. In both Sanger and Illumina workflows, bead purification scales with the number of colonies. In the conventional Illumina workflow, bead purification is performed on each reaction in parallel after the first round of a two-step PCR workflow. While our custom workflow has a higher primer cost than the Sanger workflow, the large cost reduction in other factors produces a lower overall cost than the Sanger and conventional Illumina workflows. In addition, our custom workflow has a lower time cost due to the omission of DNA prep, parallel bead purification on separate products, and secondary PCR reactions. Master mix remains a major contributor to our custom scheme’s cost, so miniaturization of reagent volumes is key to further reducing the cost of scaled microbial genotyping workflows.

In conclusion, our workflow enables scalable genotyping of microbial isolates with ∼90% success. Here, we genotyped microbes and focused on the 16S V4 region, but the workflow can be easily modified to target other genes or regions, such as the 18S region of plants or the internal transcribed spacer (ITS) region of fungi. While there is often a tradeoff between the simplicity and performance of a workflow, we have demonstrated a workflow that is relatively simple and conducive to scalability, without major sacrifices to performance. This enables efficient genotyping of large existing collections of isolates for applications such as microbial ecology, clinical microbiology, and natural product discovery.

