## Supplementary figures and images for "Scalable Genotyping of Microbial Colonies"

### Supplementary Figure S1

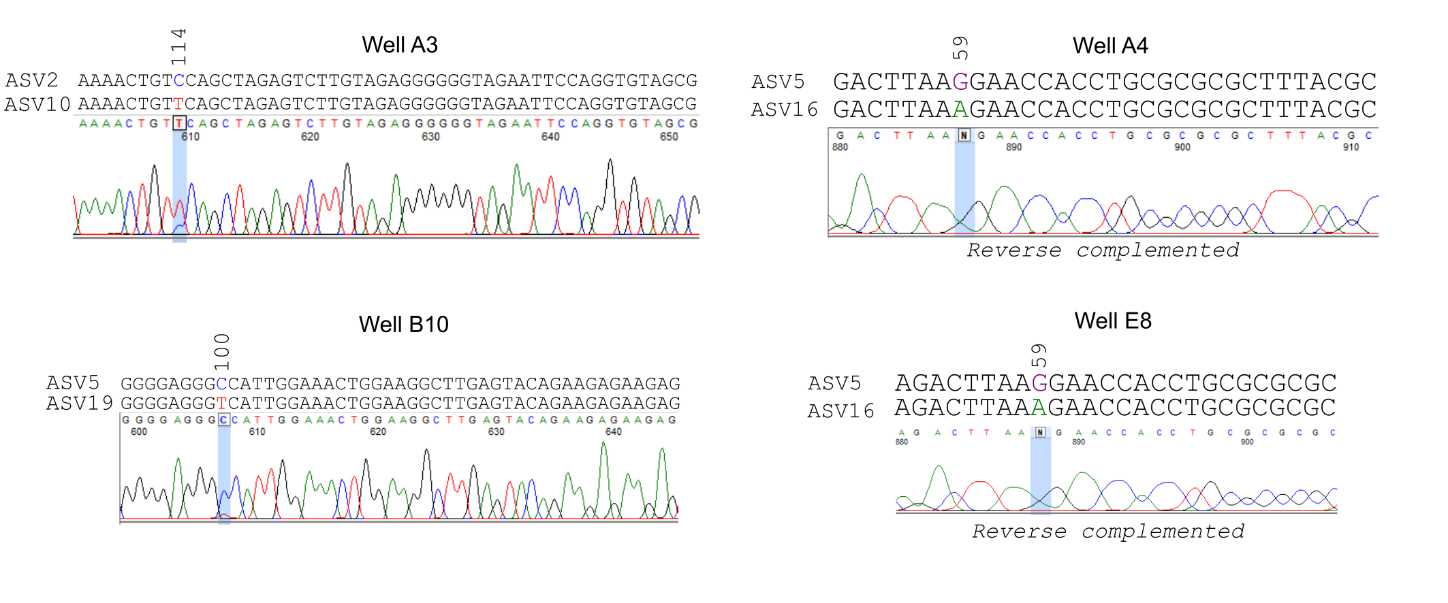
